# Integrating microscopy and transcriptomics from individual uncultured eukaryotic plankton

**DOI:** 10.1101/2024.08.25.609620

**Authors:** Catherine Gatt, Yike Xie, Kanu Wahi, Emma M. V. Johansson, Fabio Zanini

**Author notes:** Dept. of Medical Biochemistry & Cell Biology, University of Gothenburg, Gothenburg, SE-413 90, Sweden.

## Abstract

Eukaryotic plankton comprises organisms as diverse as diatoms and pelagic larvae, covering a wide spectrum of shapes, molecular compositions, and ecological functions. Plankton research is often approached using either optical methods, especially for taxonomic purposes, or genomics, which excels at describing the biochemistry of microbial communities. This technological dichotomy hampers efforts to link the morpho-optical properties of each species with its genetic and biomolecular makeup, leading to fragmented information and limited reproducibility. Methods to simultaneously acquire multimodal, i.e. optical and genetic, information on planktonic organisms would provide a connection between organismal appearance and function, improve taxonomic prediction, and strengthen ecological analysis. Here we present Ukiyo-e-Seq, an approach to generate paired optical and transcriptomic data from individual eukaryotic plankton. We performed Ukiyo-e-Seq on 66 microscopic organisms from Coogee, NSW, Australia and assembled transcriptomic contigs using a merge-split strategy. While overall phylogenetic heterogeneity spanned hundreds of taxa, diversity in individual wells was low, enabling accurate classification of both microbial plankton and marine larvae. We then combined Ukiyo-e-Seq with AlphaFold 3, a protein language model, and could confidently infer (i) the joint structure and interactions of 34 photosynthesis proteins from a single *Chaetoceros* diatom, and (ii) the cellular and developmental functions of novel proteins highly expressed in one trout larva. In summary, Ukiyo-e-Seq is a precise tool to connect morphological and genetic information of eukaryotic plankton.

## INTRODUCTION

Microscopic eukaryotic plankton inhabiting the surface layers of marine ecosystems play pivotal roles in nutrient cycling, carbon fixation, and energy transfer [1–3]. Phytoplankton are responsible for approximately half of the Earth’s primary production [3]) and a promising source of biofuel [4,5] but can also cause harmful algal blooms, with severe environmental consequences as well as human morbidity and mortality [6]. Microscopic zooplankton and the larvae of many pelagic species (fish, oysters, etc.) are also carried by oceanic currents, giving rise to a staggering biodiversity [7]. Recent estimates by meta-omics surveys [8] put the global biodiversity of eukaryotic plankton at over 150,000 distinct taxonomic units [1].

The most accurate method to characterise the heterogeneity and functions of a plankton community is to first determine its taxonomic composition and subsequently examine the molecular and biochemical capabilities *of each species of interest*. While taxonomic mapping is routinely performed via optical microscopy [9,10] and meta-omic methods such as 18s sequencing [11], the isolation and biomolecular analysis of individual species is slow, labour-intensive, and often restricted to species culturable in the laboratory [12]. Shotgun metagenomics and metatranscriptomics are excellent tools to study the biochemistry of an entire community [13] and can be combined with imaging to generate paired morphological and sequencing data at the community [8], [14] or mini-community level [15]. Nonetheless, these approaches lack resolution compared to single-organism investigations. Moreover, results can be difficult to reproduce accurately because of natural fluctuations in the taxonomic composition of each community. Single-cell genomics of environmental bacteria has matured in recent years [16,17], however eukaryotic plankton exhibit much larger variation in both genome and organismal size, hampering single-organism isolation (e.g. via traditional cell sorters) and whole-genome coverage. The ideal experimental tool would pair rapid taxonomic screening of a plankton community with high-coverage sequencing of individual organisms of interest, including unculturable ones, of any shape and size.

Here, we report the development of Ukiyo-e-Seq, an approach to generate paired multimodal (imaging and transcriptomic) data from individual, uncultured plankton organisms. We applied Ukiyo-e-Seq to 66 plankton sampled near Coogee, NSW, Australia and were able to assemble genetic contigs spanning all four superkingdoms, both public and private to a single isolate. Analysis of specific individuals enabled taxonomic identification from a combination of image and sequences. We found that some wells contained a single type of organism but others showed evidence of multiple coexisting organisms. Sequences from an individual green fluorescent phytoplankton were sufficient to reconstruct the function and three-dimensional structure of the entire photosystems I and II pathways, while novel proteins statistically related to cell cycle and embryonic development were expressed at high level in a fish egg. Overall, Ukiyo-e-Seq is a flexible approach to map extant and discover new taxonomic and molecular biodiversity at the single-organism level.

## RESULTS

### Ukiyo-e-Seq links images with transcriptomics for plankton individuals

To characterise plankton biodiversity across imaging and omics simultaneously, environmental sampling, optical microscopy, robot-assisted capture, and transcriptomics were combined into an integrated workflow, which we called Ukiyo-e-Seq (IPA: /uːˌkiːjəʊˌeɪˈsiːk/) (**Figure 1A**). “Ukiyo-e”, a style of woodblock printing art, roughly translates from Japanese as “pictures of the floating world” [18]. To demonstrate the method, 1 litre of ocean water was collected on 10/12/2021 and 21/12/2021 from Wiley’s Baths near Coogee, NSW, Australia, using a plankton net. A tea sieve and filter paper were used to remove large (>1 mm) and small (<25 μm) particles, respectively, with additional centrifugation and sedimentation steps to aid size selection (see Methods). A microscope-mounted cell picker was then used to image in brightfield and three epifluorescence channels the plankton and, using glass capillaries, transfer select organisms into wells of a microtiter plate. Libraries for single-well RNA sequencing were prepared using Smart-seq2 adapted to 384-well plates [19,20]. In total, 66 wells were imaged and sequenced, plus 4 negative control wells.

**Figure 1:**
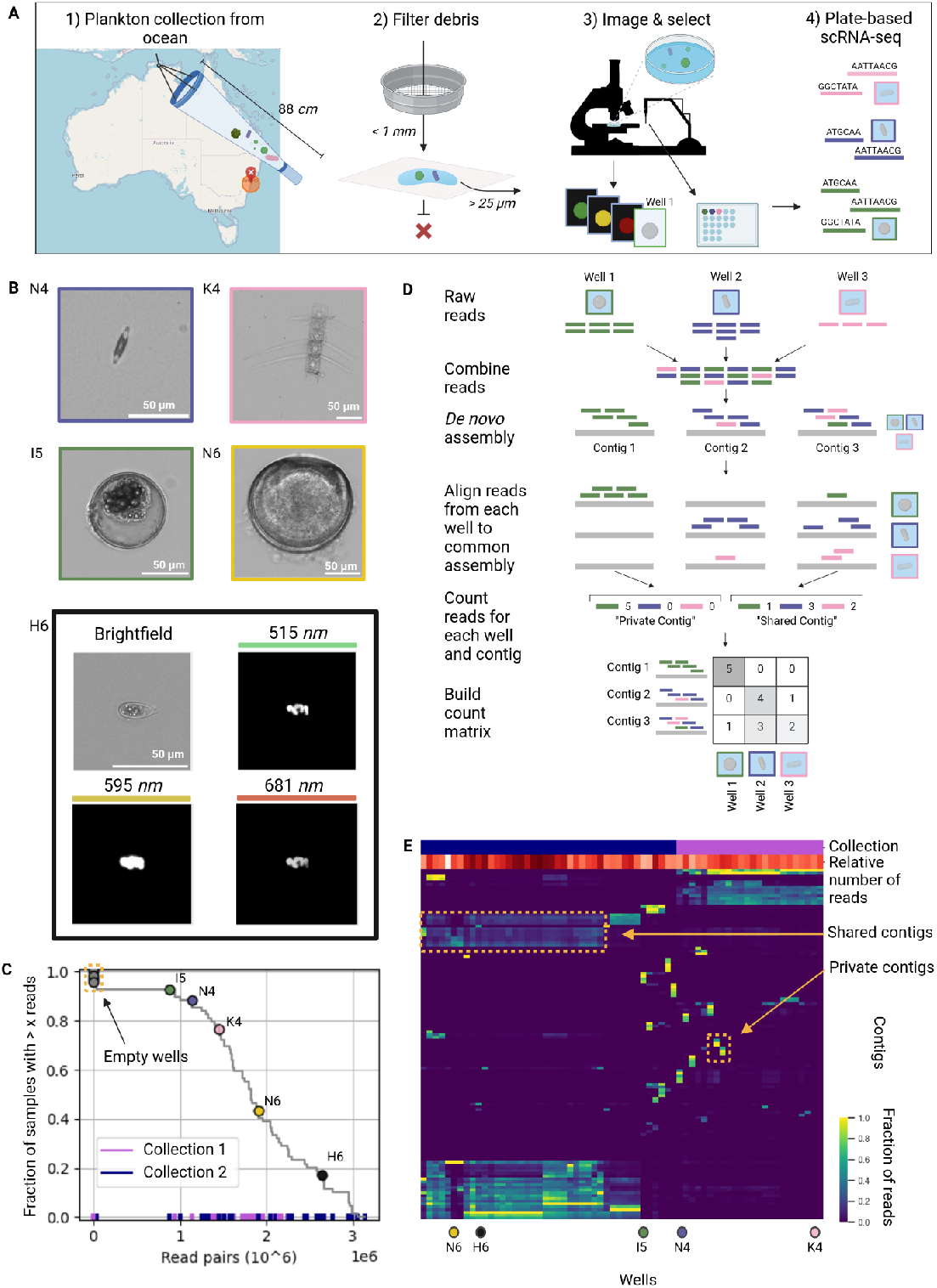
Ukiyo-e-Seq collects paired imaging and transcriptomic data from individual uni- and oligocellular marine organisms. **A**. Overview of Ukiyo-e-Seq, starting from organismal enrichment with a plankton net (1, keep particulate) and later tea sieves and filter paper (2, keep middle fraction), followed by image-assisted picking (3) and single-well RNA-Seq (4). **B**. Images for 4 representative wells in brightfield (middle) and 1 additional representative well in brightfield and epifluorescence (bottom, emission wavelengths shown) for samples acquired near Coogee beach, NSW, Australia **C**. Cumulative distribution of sequencing coverage across 70 wells in two separate experiments. Four empty wells were used as negative controls. **D**. Schematic of the merge-split metatranscriptome assembly pipeline. **E**. Read coverage heatmap across organisms of the most abundant assembled contigs, showing both shared/public contigs (blocks) and private ones (vertical stripes).

Because individuals could be selected for picking one at a time, a morphologically diverse library of organisms was collected (**Figure 1B and Figure 1 - Supplementary 1**). Quality controls on the raw sequencing data showed that 90% of samples yielded more than 1 million read pairs and 40% more than 2 million (**Figure 1C**). The only 4 samples with less than 10,000 read pairs were the negative controls (255, 453, 509, and 6231 read pairs). No clear relationship was observed between sequencing coverage and optical properties including organismal size (**Figure 1B, middle**) and fluorescence intensity (**Figure 1B, bottom**).

### Metatranscriptome assembly of planktonic reads via a merge-split strategy

Reference genomes are not available for most oceanic species. Therefore, the transcriptomic reads were assembled *de novo* using rnaSpades (Bushmanova et al., 2019). To mitigate the effects of low per-sample coverage, a merge-split strategy was developed (**Figure 1D)**: (i) Reads from all wells were pooled and assembled to form a common reference, then (ii) Reads from each well were aligned against this reference separately. A total of 125 million paired end reads and 4 million unpaired reads (see Methods) were assembled into 801,126 contigs with a maximum contig length of 114,708 bases, a total length of 273,350,101 bases, L50 = 474 bases and N50 = 204,309 bases. The number of reads from each sample that align to each contig were compiled into a single matrix [21].

### Planktonic contigs can be public or private

To gain broad biological insight from the data, the submatrix with the top five most highly expressed contigs from each well was visualised (**Figure 1E**). Two main patterns of expression were observed. First, rectangular blocks of high-coverage contigs, spanning most samples from one single collection day, were detected. Because plankton’s water was thoroughly exchanged with distilled water before picking, these contigs suggest that many plankton might be covered with “genetic dust” that is specific to any given day. Alternatively, they could derive from experimental contamination during the sample processing, although that would not explain why the contigs from the two biological replicates are so different between each other. Private contigs, highly expressed in one well only, were also observed among the top 5 most expressed contigs in some wells. These well-specific contigs likely originated from the organism under the microscope, i.e. could be linked back to the optical properties (e.g. shape, colour) of each planktonic individual. No relationship was observed between the presence of highly abundant private contigs in a well and its sequencing depth (**Figure 1 - Supplementary 2**).

### Single-well taxonomic sequence analysis covers four superkingdoms

We then used Kraken 2 to compare all contigs against a large database of metagenomic and metatranscriptomic data and thereby understand the taxonomic identity of public and private contigs despite the lack of reference genomes (**Figure 2A**) [22]. The taxonomic depth of these predictions decreased if either the similarity to known sequences was low or if the matching database sequences had themselves a shallow taxonomic assignment. Across all samples, contigs were found to be extremely heterogeneous, covering all four superkingdoms (eukaryota, bacteria, archea, and viruses, see **Figure 2 - Supplementary 1**). Of all reads, 54% were classified as eukaryotic, 13% as bacterial, 0.30% as viral, 0.15% as archaeal, and 33% unknown or unclassified. The union of the 20 most abundant taxonomic units in each well contained hundreds of taxa, including high abundance of marine microorganisms such as *Calanus* and macroscopic organisms such as *Bivalva*. Only 5 of the 50 most abundant taxa across all samples were not marine organisms: *Bradyrhizobium sp. PSBB068* (a soil bacterium), *Homo sapiens (*human*), Vitis rotundifolia* (a grapevine), *Hordeum vulgare* (common barley) and *Endopterygota* (a winged insect). These could be laboratory contaminants or be present in coastal waters near a densely populated shoreline.

**Figure 2:**
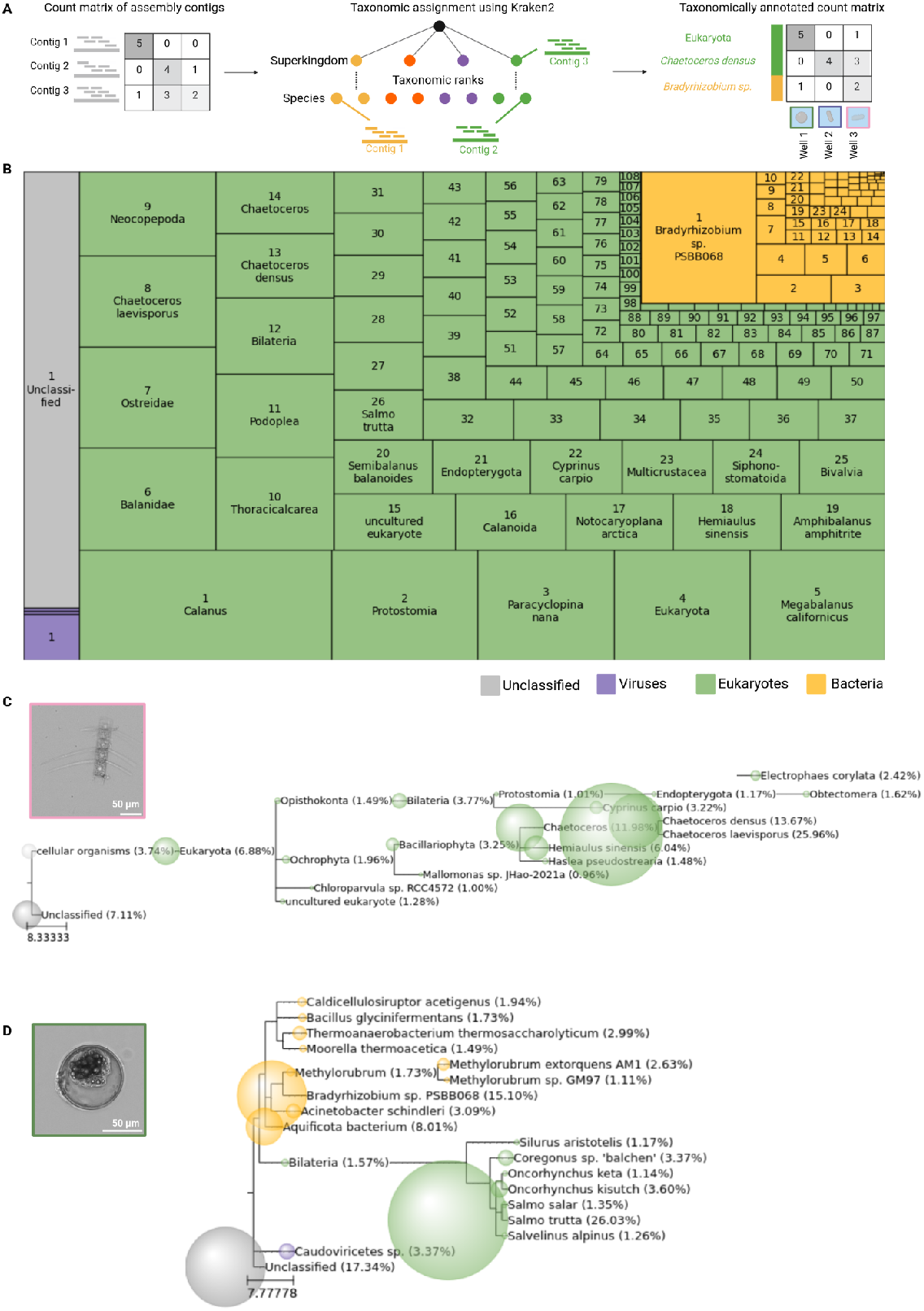
Global and single-well taxonomic analysis of plankton transcriptomes. **A**. Analytical workflow to compute taxonomic coverage maps for each well. **B**. Abundance treemap of taxonomic units. Taxa are numbered within each superkingdom by the number of reads mapping to that taxonomic unit (lower numbers indicate larger read counts). **C, D:** Paired images and phylogenetic trees on sequence contigs for two representative wells, containing a photosynthetic diatom (**C**) and a brown trout egg or larva (**D**). The size of each ball is proportional to the fraction of contigs assigned to that taxon by Kraken 2. Colours by superkingdom (green: eukaryotes, orange: bacteria, purple: viruses, grey: unassigned).

### Single-well phylogenies clarify taxonomy and suggest patterns of coexistence

Despite the high degree of biodiversity found across samples, individual wells were significantly less heterogeneous and taxonomically sharply defined. The 20 most abundant sequences from well K4 were almost exclusively assigned to the *Chaetoceros* genus, a type of photosynthetic diatom. This prediction was also supported by the organism’s unique morphology (**Figure 2C**). Patterns of potential coexistence were also observed. Well I5 contained 26% of reads assigned to *Salmo trutta* (brown trout) which, together with the morphology from the microscopy image, indicated that the main organism was a fish egg or early larva. However, the same well also contained a few bacterial species that were not observed in all wells (**Figure 2D**). Overall, these results demonstrated that the multimodal data with single well resolution generated using Ukiyo-e-Seq enabled clear taxonomic assignment of individual organisms, overcoming the limitations of both microscopy and metagenomics alone.

### Reconstruction of protein complexes from a single environmental organism

Next, we assessed the utility of Ukiyo-e-Seq for functional characterisation of environmental organisms. Based on morphology, taxonomy, and epifluorescence data, multiple organisms were likely to possess photosynthetic capabilities. To confirm photosynthetic activity in specific wells, we identified open reading frames (ORFs) in the contigs expressed by each well using the NCBI ORF finder [23], then functionally annotated ORFs [24] and extracted all ORFs assigned to protein members of the KEGG photosystem I and II complexes (**Figure 3A**). Multiple wells, such as K4 and N4, contained almost all members of both complexes, further buttressing their taxonomic assignment, while other wells that were assigned to non-photosynthesising taxa contained no or only a few detectable ORFs in this pathway (**Figure 3B**). As an illustration of these results, we extracted an ORF from well K4 annotated as photosystem II protein D1 (psbA) and compared it against recently reported sequences from related planktonic taxa (**Figure 3C**) [25]. The alignment showed an especially high similarity with 3 out of 7 species, with 340 residues sharing 100% sequence identity across all four protein sequences. We then used AlphaFold 3, a large language model (LLM) for protein folding [26], to predict the three-dimensional structures of the photosystem II (**Figure 3D**) and I (**Figure 3 - Supplementary 1**) complexes of the *Chaetoceros* individual from well K4. *In silico* photosystem II folding covered 22 peptide chains and 3,620 residues, with 76% of the atoms assessed to a level of “confident” or above (**Figure 3D, inset**), while photosystem I covered 12 proteins and 2,586 residues (84% atoms “confident” or above). Taken together, both complexes comprised 34 chains and 6,206 residues (80% confidently predicted), all from the same microorganism.

**Figure 3:**
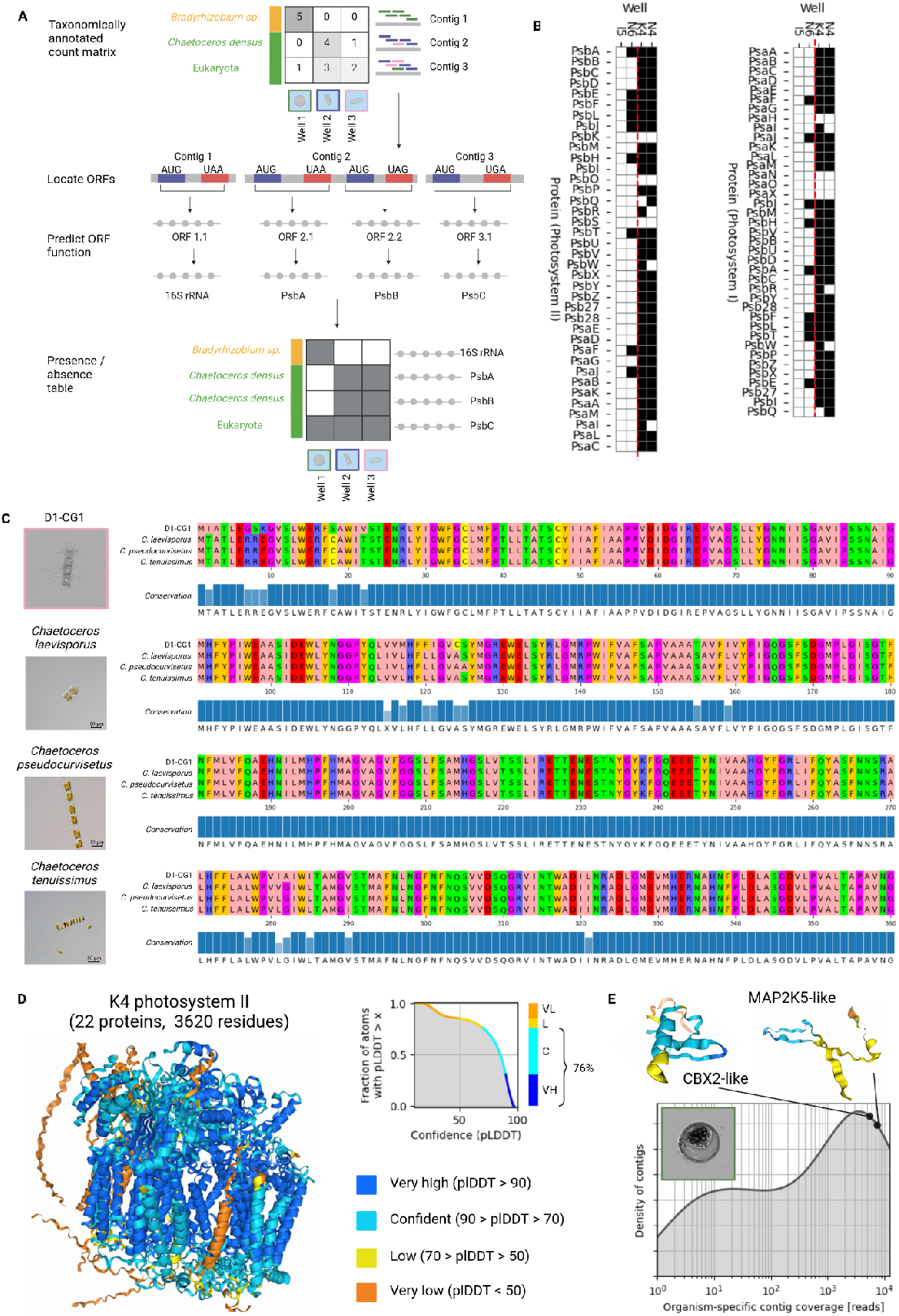
Functional and structural analysis of proteins and complexes from individual organisms. **A**. Workflow of functional analysis of proteins found in individual wells. **B**. Presence of genes belonging to the KEGG photosynthesis pathway across four representative wells. **C**. Multiple sequence alignment of a newly identified D1-CG1 photosystem II reaction center protein D1 (PsbA) sequence identified in well K4 with three other PsbA sequences and corresponding pictures from [25]. **D**. Three-dimensional structural reconstruction of photosystem II from well K4. **E**. Distribution of expression for contigs assigned to *Salmo trutta* in well I5 (image in top left inset) and structural prediction of two predicted proteins from highly expressed contigs. Structural predictions via AlphaFold 3 [26].

Finally, we evaluated the ability of Ukiyo-e-Seq to gain genetic and functional insight into early-stage development of larger marine organisms by focusing on well I5, previously determined to contain an egg or early embryo of *Salmo trutta*. Sequence and structural analyses of *Salmo trutta* ORFs in highly expressed contigs identified novel proteins with sequence and structure relationships to known members of the cell cycle and embryo development pathways, including chromobox 2 (*CBX2*, 5485 reads / contig) and dual specificity mitogen-activated protein kinase kinase 5 (MAP2K5, 7377 reads / contig) (**Figure 3E**). Importantly, no photosynthesis genes were detected in this well (**Figure 3B**), indicating a low rate of false positives for functions that are not actually present in the organism of interest. Overall, these data demonstrated that Ukiyo-e-Seq enabled the functional and structural characterisation of novel proteins and complexes starting from a single, uncultured organism, providing a biological link between organismal taxonomy, morphology, genetics, and behaviour.

## DISCUSSION

Eukaryotic plankton play essential roles in the Earth’s biotic and chemical cycles: atmospheric oxygenation, primary production, and the base of the food pyramid, among others [27]. These functions are carried out by a spectrum of organisms highly diverse in both appearance and genetics. For instance, dozens of species within the *Chaetoceros* genus alone cohabit the Jiaozhou bay near Qingdao (China), each with its unique genetic signature, morphology and, presumably, ecological function [REF]. To decipher the complex balance of species found in most aquatic environments, technologies able to characterise not only whole communities but the individual plankton within are needed. The method presented in this study, Ukiyo-e-Seq, fills this gap by achieving multimodal characterisation of single planktonic organisms of choice: each image is accompanied by around a million transcriptomic reads.

Ukiyo-e-Seq has distinct advantages over traditional approaches such as microscopy alone (e.g. diatoms.org [9]). First, image and sequences can be matched against separate reference databases to increase the robustness of taxonomic assignment (**Figure 2C**). This could also be useful to unveil organismal (e.g. Batesian) mimicry [28]. Second, open reading frames can be extracted from each organism’s transcriptome to identify highly expressed proteins and biochemical pathways (**Figure 3B**), establishing a direct link between an organism’s appearance and its molecular content. Given the low taxonomic heterogeneity within each well (**Figure 2C-D**), Ukiyo-e-Seq leads to more interpretable results than meta-omics and mini-metagenomics [15], which lack single-organism resolution. Third, compared to valve-[15] and droplet-based microfluidic platforms [16], Ukiyo-e-Seq can profile a wider range of organismal sizes (up to hundreds of microns) because glass capillaries with different bore sizes can be used in the same experiment. This greatly expands the types of eukaryotic plankton that can be examined. A disadvantage of Ukiyo-e-Seq, compared to standard community-level approaches, is the requirement for specific training on the cell picker. This makes Ukiyo-e-Seq better suited for deep investigations than routine monitoring [27].

One unique application of Ukiyo-e-Seq is the recovery of multiple expressed protein sequences from the exact same organism. Coupled with deep learning models [26][29], this enables the reconstruction of three-dimensional structures of entire protein complexes, including interchain interactions, starting from a single, uncultured plankton (**Figure 3D**). This is an important improvement over current methods which require culturing [30] [31], are limited to one protein at a time [32], or specific to bacteria [16]. Given the current pace of artificial intelligence development, it might become possible to discover entirely new biochemical pathways from an individual plankton, sampled directly from the environment.

In conclusion, Ukiyo-e-Seq enables the collection of paired imaging and sequencing data from individual eukaryotic plankton of choice, making it ideally positioned to map marine biodiversity and accelerate the discovery of novel proteins and complexes from environmental samples.

## ACKNOWLEDGEMENTS

We would like to thank Annalice Creighton and the Australian National Maritime Museum for lending us their plankton net and Rob Salomon at Children’s Cancer Institute for gracefully agreeing to let us access their robotic pipettor. We would also like to thank the late Leanne Armand, Linda Armbrecht, Justin Seymour, and Harriet Alexander for scientific discussions and feedback on early versions of the manuscript.

## METHODS

### Plankton collection

Surface-water samples were collected at Coogee beach, NSW (33°55’32.4”S 151°15’32.9”E) with the use of a plankton net (Australian Entomological Supplies, E50F Plankton Net with Collecting Bottle), with a mesh size of 100 μm, 300mm in diameter, and net 0.88 metre long, which was kindly lent by the Australian National Maritime Museum. Samples were collected on two separate occasions (10/12/2021 and 21/12/2021). On each occasion, two 1.5 L bottles of seawater were collected, one from the surf near the beach and one from the adjacent rock pool (Wiley’s Baths). The pool’s water is in constant exchange with the open ocean.

### Sample Preparation

For each collected bottle, seawater was passed through a tea strainer (Woolworths Ltd mint tea strainer, mesh size ∼1 mm) to remove large debris into an autoclaved glass beaker, then left to settle for 30 minutes on the bench at room temperature. 50 ml of supernatant were transferred into a Falcon tube, avoiding the sediment at the bottom, and centrifuged at 3,500 g for 10 minutes to concentrate the plankton at the bottom. The supernatant was discarded, and the pellet was resuspended in distilled water. The sample was then poured onto 25 μm filter paper, discarding the flowthrough and later washing the paper runoff into a 6-well flat-bottom tissue culture plate.

### Cell Picking

Cells were captured using the CellCelector platform (ALS Automation GmbH/Sartorius) with a glass capillary tube (inner diameter: 150 μm) using the manual picking mode. The cells were dispensed into a 384-well Frame-Star plate (4titude Ltd, Surrey, UK) containing 1 μL lysis buffer, which was cooled to 4 ºC for the duration of the capture. For the first batch of samples, a brightfield image of each cell was recorded before capture. For the second batch of samples, brightfield and fluorescence images (emission wavelengths: 515, 595 and 681 *nm*) of each cell were taken before each capture. For both batches, a brightfield image was taken after each cell capture to confirm successful picking. Two of the wells included in the following cDNA synthesis, library preparation and sequencing did not have cells placed into them, to act as a control for background contamination originating from the laboratory equipment used. Following cell capture, the lysis plate was stored at –80 C for one month.

### cDNA synthesis and library preparation

cDNA synthesis was performed following the Smart-seq2 protocol [33]. In brief, 1.8μl of reaction mix (SmartScribe, Rnase Inhibitor, 5x First Strand Buffer, DDT (100mM), Betaine (5M), MgCl2 (1M), TSO (100uM), H2O) was added to each well containing a single cell lysate using the MANTIS automated liquid handler and reverse transcription was performed by incubating the plate at 42 C for 90 minutes, then 70 C for 5 minutes using a thermal cycler (check brand). Following this, 4.2μl of PCR mix (H2O, Kapa Buffer (5x), dNTP (10mM), Polymerase (1U/ul), IS_PCR (10uM)) was added to each well using the MANTIS automated liquid handler (brand) and PCR was performed according to the following cycle: 1) 95 C for 3 minutes, 2) 26 cycles of 98 C for 20 seconds, 67 C for 15 seconds, 72 C for 5 minutes, and 3) 72 C for 5 minutes. A mean concentration of the cDNA in each well was calculated using measurements from the Bioanalyzer 2100 (Agilent Technologies, Waldbronn, Germany) and cDNA concentrations were normalised to 1.4 ng/μl with nuclease-free water using the automated liquid handler Mosquito. The plate was stored at 4C until library preparation. The Nextera XT kit (Illumina, San Diego, CA, USA) was used for tagmentation using 15 cycles of PCR. Bead purification was carried out using Agencourt Ampure XP (Beckman Coulter, Brea CA, USA) magnetic beads. One sample was purified at a ratio of 0.7X for two cycles and a second sample was purified using an additional reverse purification at a ratio of 0.5X for one cycle. Library concentration and size were compared between the two samples using the High-Sensitivity D1K TapeStation (Agilent 2200 TapeStation).

### Sequencing

Libraries were sequenced on an Illumina NextSeq 500 (Illumina, San Diego, CA, USA) using a reagent v3 kit (2×300 bases), at a depth of about 250,000 read pairs per well at Ramaciotti Centre for Genomics (UNSW).

### Quality Control

Reads were filtered using Trimmomatic version 0.39 [34] using paired end mode, allowing 2 mismatches requiring 30 bp of overlap between R1 and R2 to identify the fragment as being less than the read length and requiring 10bp of sequence to match before removing the read to filter out low quality reads and the Illumina adapter sequences. Reads containing the template switch oligo (TSO) sequence (AAGCAGTGGTATCAACGCAGAGTACATGGG), the reverse complement of the TSO sequence, and the sequencing artefact (30G) were removed using a custom python script, retaining both unpaired and paired reads. Total read counts for each well were calculated by adding the number of paired reads, unpaired forwards and unpaired reverse reads that passed the above quality assessment. A cumulative distribution function of the total read across individual wells was generated using Python 3.11.

### Contig Construction

Paired forwards and reverse reads from all wells were combined into two single files and these combined reads were assembled into contigs using rnaspades (v3.15.5) with default parameters [35]. Bowtie2 [36] was used to map all the paired and unpaired reads of each well individually against the contigs assembled above. Then, pileup.sh from BBMap (v38.18) [21] was run on individual bam files to count the number of forward and reverse reads from each individual well that map to each contig . An expression matrix of all contigs in the combined rnaSpades assembly, and the total number of reads (forwards plus reverse) from each well mapping to that contig was generated. The top five contigs with the highest number of reads mapped to them from each well were visualised in Python 3.11.6 using double hierarchical clustering.

### Taxonomic Annotation

For taxonomic classification, Kraken2 2.0.7-beta [22] was used with the NCBI non-redundant nucleotide database as a reference library with default parameters to assign a taxonomic ID to each of the contigs. The number of reads that mapped to each contig that was assigned to the same taxonomic ID by kraken were summed together to create a taxonomically annotated count matrix.

The contigs which were given a taxonomic ID 1 or 0 were assigned as ‘unclassified’. All other taxonomic IDs other than 1 or 0 were assigned to a ‘superkingdom’ (Eukaryota, Bacteria, Viruses and Archaea) using the ETE 3 package [37]. The total number of reads mapped to each of the taxonomic IDs assigned to each superkingdom or unclassified group across all wells were added up and divided by the total number of reads mapped to all taxonomic IDs to calculate the percentage of each group (Eukaryota, Bacteria, Viruses, Archaea and Unclassified). The union of the top twenty most highly expressed taxonomic IDs from each well (taxonomic IDs with the highest number of reads assigned to them) were visualised in Python 3.11.6 using squarify. The top twenty most highly expressed taxonomic IDs (taxonomic IDs with the highest number of reads assigned to them) from the individual wells ‘I5’ and ‘K4’ were retrieved and a NCBI taxonomy tree was constructed and visualised using the ETE 3 package [37].

### Functional Annotation

The common assembly contigs were assigned to each individual well if they had at least one read mapped to them from that well. Open reading frames (ORFs) were located within each contig using the NCBI ORF finder v0.4.3 [23], using the standard genetic code (transl_table=1) and default parameters. InterProScan v5.64-96.0 [24] was used using default parameters to functionally annotate the ORFs. The functionally annotated ORFs in wells ‘I5’, ‘N6’, ‘K4’ and ‘N4’ were searched for genes present in the KEGG photosystem I and II reference pathways. A presence / absence matrix of the genes from photosystem I and II across the wells ‘I5’, ‘N6’, ‘K4’ and ‘N4’ was visualised using Python. The sequence ‘D1-CG1’ photosystem II reaction centre protein D1 (PsbA) sequence identified in well K4 was aligned with three other PsbA sequences using clustal omega (1.2.4) in the job dispatcher at EMBL-EBI [38] and was visualised in Python using pyMSAviz. One sequence was selected for each gene that was present in the photosystem II of well ‘K4’ and the protein structure of the ORFs were predicted using AlphaFold 3 [26] online server with default parameters. A cumulative distribution function of the pLDDT confidence scores of each atom in the model of the structure and was visualised using Python, maintaining the same colour scheme for confidence scores as AlphaFold 3. The density of the number of reads that mapped to each of the contigs that were taxonomically assigned to ‘Salmo trutta’ in well ‘I5’ was computed and visualised using Python. Functional annotation of two ORFs, I5-FZ1 and I5-YX1 was carried out using blastp and the reference proteins (refseq_proteins) database . The structures of these ORFs were also predicted using the AlphaFold 3 [26] online server with default parameters.

### Data and Code Availability

All images and sequencing reads are available on GEO (GSE274796). The code used to produce the results is on GitHub at https://github.com/fabilab/ukiyo-e-seq.

## Supplementary Materials

**Figure S1 - Supplementary 1:**
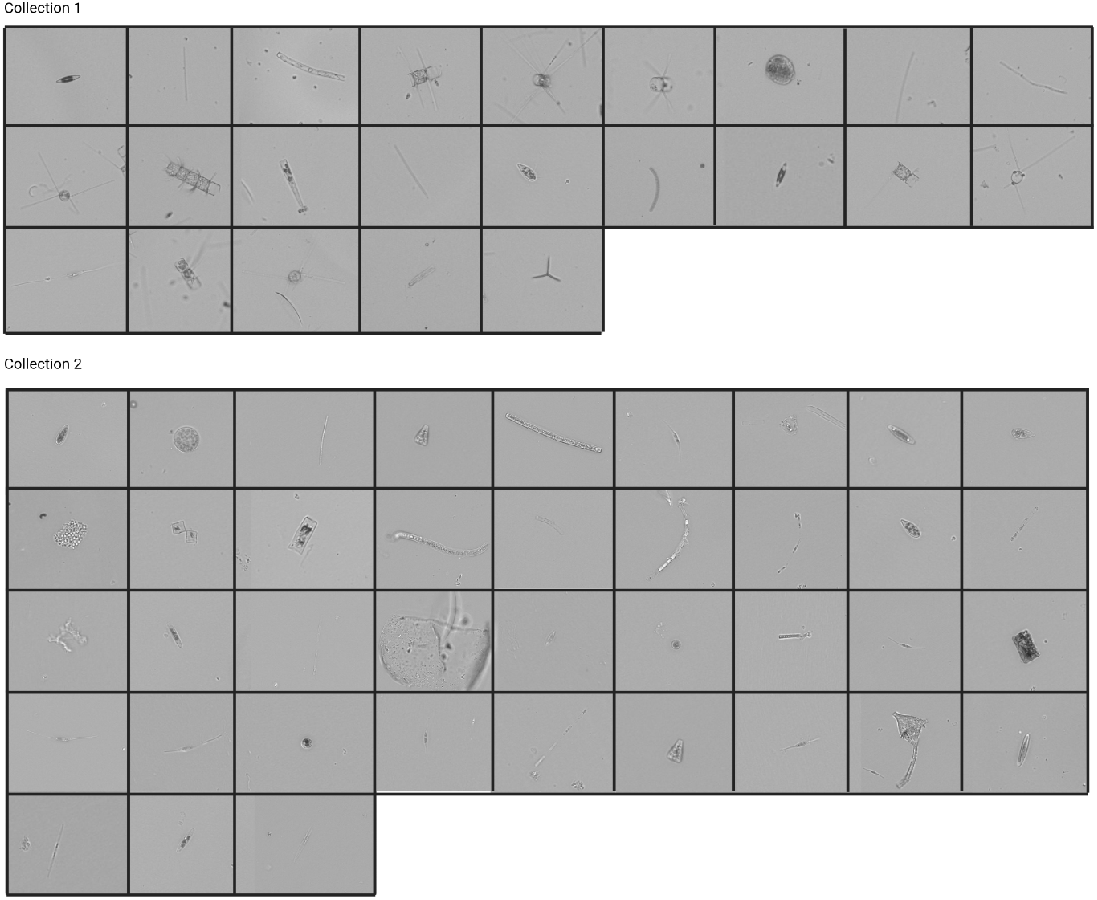
Brightfield images (zoom in) of the plankton captured in each of the 66 wells during our two experimental batches.

**Figure S1 - Supplementary 2:**
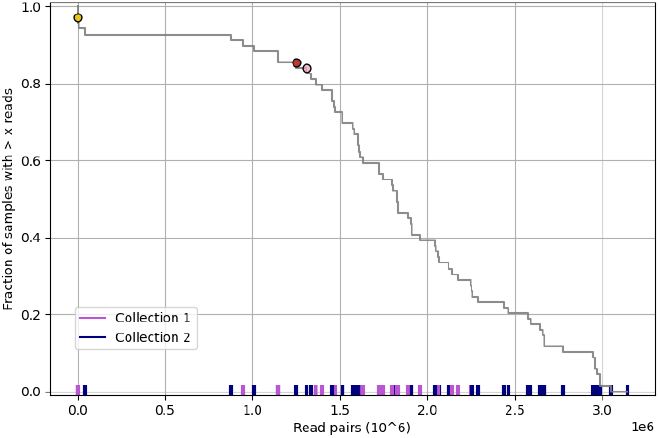
Examples of wells with private contigs. No relationship between the presence of private contigs and sequencing depth was found.

**Figure S2 - Supplementary 1:**
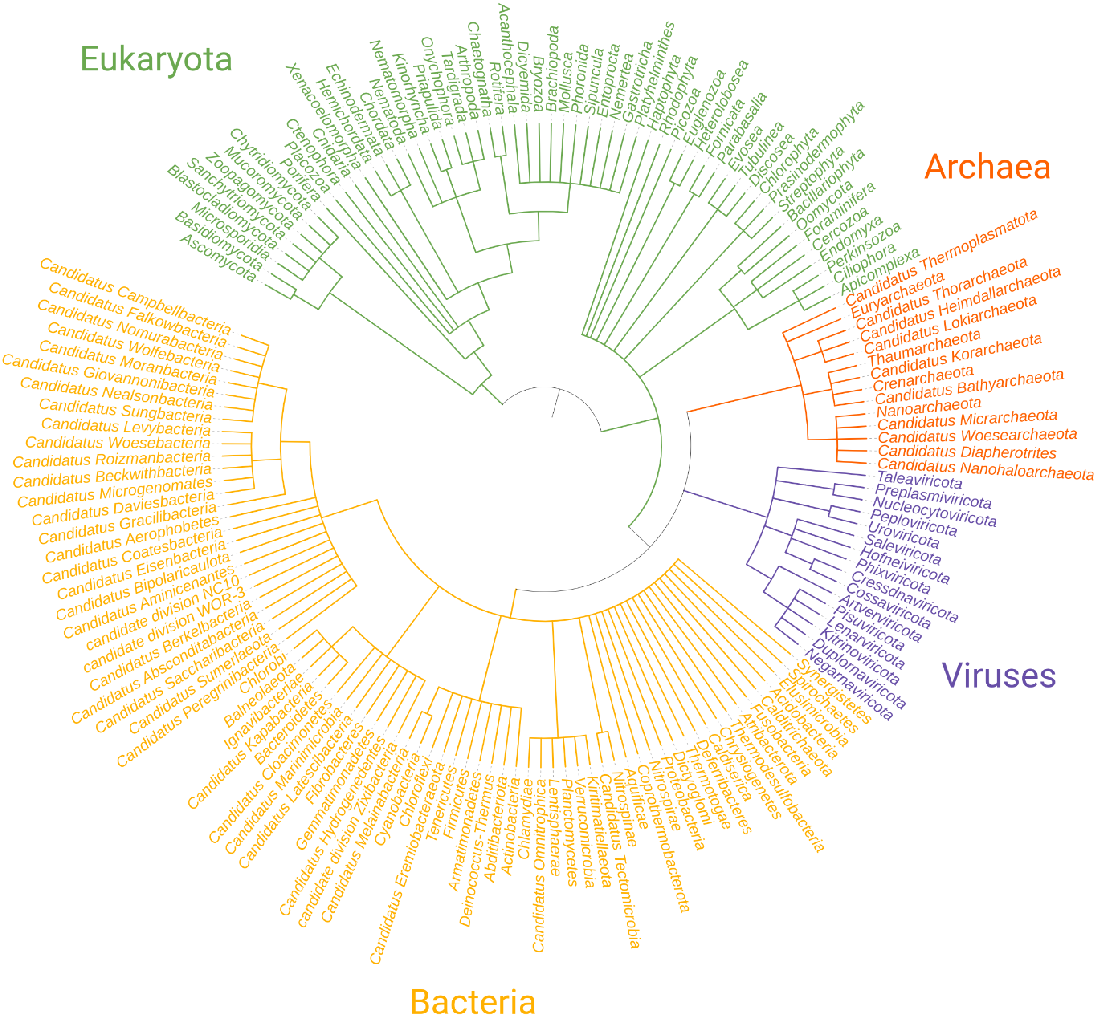
Phylogenetic tree of all taxa identified by Kraken 2 across all contigs (all wells together). Branches were collapsed at the phylum level for simplicity and coloured by superkingdom.

**Figure S3 - Supplementary 1:**
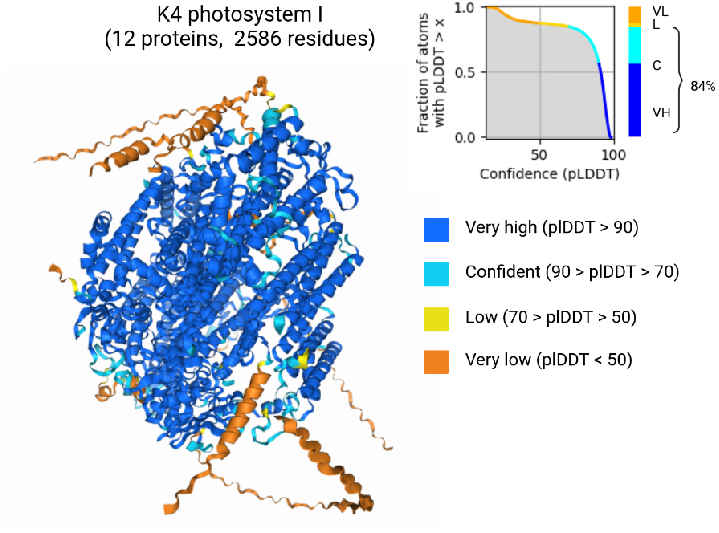
Structural reconstruction of Photosystem I from well K4, the same as in **Figure 3D**.

